# Do sex differences in autosomal recombination rates facilitate divergence?

**DOI:** 10.64898/2026.04.27.721057

**Authors:** Amanda Hansson, Marina Rafajlović

## Abstract

Recombination rate varies within and between individuals. One form of such variations is seen between sexes in dioecious populations, with males typically exhibiting a smaller recombination rate than females. This is true both for sex chromosomes and autosomes (so-called heterochiasmy). Although a large body of theory exists on the role of sex chromosomes in adaptation and population divergence, much less is known about the role of heterochiasmy. Recently, it has been suggested that heterochiasmy can facilitate local adaptation and divergence, but if, and when this is true has not been systematically studied theoretically to date. Here we use Individual-based simulations to assess the effect of sex differences in autosomal recombination rates on the process of divergence and adaptation in populations subject to divergent selection and migration. We found evidence supporting that sex differences in autosomal recombination rate between adaptive loci can facilitate, and especially maintain, divergence, but this is true only under very limited conditions, involving strong selection, high sex-averaged effective recombination rates and relatively high rates of migration compared to the strength of selection. We further found that this effect, when present, is typically weak but is amplified in cases of highly polygenic adaptation in comparison to cases with a few adaptive loci of strong effect. We conclude that, in most cases, sex differences in autosomal recombination rate alone are unlikely to noticeably contribute to the process of adaptation and divergence. Further studies are needed to evaluate their effect in combination with other processes not considered in the present study, such as assortative mating between the alike mates, or recombination suppression in heterozygotes.

**Teaser:** In dioecious populations, recombination rate typically differs between males and females. This is true both for sex chromosomes and autosomes. While much theoretical research has focused on understanding how recombination rate differences in sex chromosomes shape local adaptation and divergence, we lack theoretical knowledge of the potential role of sex differences in autosomal recombination rates. Recombination has a dual role in local adaptation. Strong recombination can effectively purge deleterious alleles, but it can also break apart beneficial allele complexes (and vice versa for weak recombination). Thus, one may expect that in the presence of both strong and weak recombination exhibited by females, and males, respectively, population divergence can be efficiently facilitated. But is this true? Here, we study this question theoretically using computer simulations. Our main finding is that sex differences in autosomal recombination can facilitate divergence, but this effect is typically weak and present only under very stringent conditions.

## Introduction

Population divergence and speciation are generally gradual and often diffuse processes governed by a complex interplay of evolutionary mechanisms, wherein some act to increase divergence and adaptation and others act to decrease it. In the context of interconnected populations evolving under different selection pressures, selection works to increase both divergence and local adaptation. This effect is counteracted by migration (Bulmer, 1972) and recombination (Felsenstein, 1981) that allow for gene flow between the populations (Felsenstein, 1981; Barton & Bengtsson, 1986; Yeaman & Otto, 2011; Yeaman & Whitlock, 2011; Eriksson & Rafajlović, 2021) as well as by drift (Rafajlović et al. 2016). In general, divergence can only be established and maintained if overall selection is sufficiently strong in comparison to the joint effect of migration, recombination and drift, such that gene flow between the populations is sufficiently small (Barton & Bengtsson, 1986).

In the process of adaptation, recombination has a dual role. Strong recombination can hinder divergence and adaptation by breaking up locally adapted allele complexes (Felsenstein, 1981). However, strong recombination can also enable the purging of deleterious alleles and combining of beneficial ones, allowing selection to act more efficiently and thereby facilitate local adaptation (Felsenstein, 1981). The opposite is true for weak recombination. A trade-off between the negative and positive effects of recombination between adaptive loci may result in an intermediate value of the recombination rate as an optimum for successful divergence (Aeschbacher & Bürger, 2014) Previous research has shown that recombination rates vary greatly between species, populations, individuals, and genomic regions (Martin et al., 2019; Venu et al., 2024). Existing theory has mainly studied the role of differences in recombination rates on sex chromosomes as a special case of the recombination-rate variation between individuals (Haldane, 1922; Charlesworth et al., 1987; Fraïsse & Sachdeva, 2021).

In species with sex chromosomes, recombination does not occur between sex chromosomes in the heterogametic sex (between X and Y in mammals and W and Z in birds, for example) and the X (or Z) chromosome often experiences reduced recombination rates compared to autosomes (Fraïsse & Sachdeva, 2021). It has been shown that these differences may be important drivers of divergence (Fraïsse & Sachdeva, 2021). Much theoretical and empirical work has been dedicated to providing explanations of how this difference influences divergence and adaptation, including, but not limited to, Haldane’s rule (Haldane, 1922), or the faster-X theory (Charlesworth et al., 1987; Charlesworth et al., 2018).

However, in addition to the differences in recombination rates between sex chromosomes, previous research has shown that differences in autosomal recombination rates between males and females are present in many species (Lenormand & Dutheil, 2005), and cases exist where autosomal recombination is completely absent in the heterogametic sex (Kong et al., 2010). Although several hypotheses have been proposed as to why sex-differences in autosomal recombination rates may have evolved (Barboza & Blackmon, 2025), the evolutionary implications of this phenomenon remain to be understood.

In a recent study, Venu et al. (2024) explored this question using a three-locus model of divergence tailored to mimic divergence of three-spined sticklebacks in a system of marine and freshwater habitats.

They concluded that differences in autosomal recombination rate between males and females facilitate divergence.

However, Venu et al. (2024) considered only specific conditions in their models, such as very strong selection and comparatively strong migration. In addition, their model included sex differences in migration rate. Interestingly, the strongest effect of the recombination rate differences between the sexes was obtained in their model variant that further included recombination suppression in cis (or coupled)-heterozygotes (i.e., recombination is suppressed for heterozygotes that have locally beneficial alleles linked on one homologous chromosome, and locally deleterious alleles linked on the other homologous chromosome). Although these conditions are suitable for their study system, the question remains whether their findings are a rule or an exception. In the present study, we ask: Can sex differences in autosomal recombination rate alone facilitate the process of divergence between populations subject to different environmental conditions? If yes, under which conditions?

To answer these questions, we use computer simulations of population divergence wherein sex differences are allowed only in recombination rate between adaptive loci on autosomes. Our model further extends the model by Venu et al. (2024) by assuming polygenic basis of adaptation, an assumption supported by empirical evidence that adaptation in many species is polygenic (Marques et al., 2019). We assess model outcomes over a wide range of values of model parameters, including selection strength, migration, sex-average autosomal recombination rate, and a parameter governing the extent of male-female differences in autosomal recombination rate.

Our main finding is that the differences in autosomal recombination rate between the sexes can facilitate divergence, or contribute to its maintenance, but only under very limited conditions, such as very strong selection, migration and recombination.

## Methods

To assess the potential effects of differences in recombination rate between males and females on the process of divergence and adaptation, a stochastic forward-in-time multi-locus model was constructed. We model two diploid populations subject to migration, fecundity selection, recombination, mutation, and locally random mating. The model was realised by means of computer simulations written in the Julia programming language (Bezanson et al., 2017). Data analyses and visualisation were performed using the R programming language (R Core Team, 2025) and the R package ggplot2 (Wickham, 2016).

### Model description

For simplicity, the model uses two demes of equal and fixed diploid size (*N*), equal sex ratios and with equal per-individual per-generation migration probabilities between the two demes. Generations are non-overlapping. We assume polygenic adaptation with *L* adaptive loci. For simplicity, all loci are assumed to be bi-allelic, with the two alleles denoted by *A* and *a*.

The order of simulation steps in each generation is: migration, selection, gamete production and recombination, mutation, and mating.

Migration is implemented by sampling uniformly at random (without replacement) emigrating individuals from each deme with a per-individual migration probability *m*. In our simulations (except for Scenario 3, see Appendix A), *m* is chosen in relation to the per-allele selection coefficient (*s*), i.e, 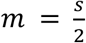 *m* = *s*, or 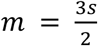 (Table 1). These values correspond respectively to the weak, intermediate and strong migration in comparison to selection (Bulmer, 1972; Yeaman & Otto, 2011). The emigrants are transferred to their nonnative deme. This is followed by fecundity selection and gamete production.

**Table 1.**
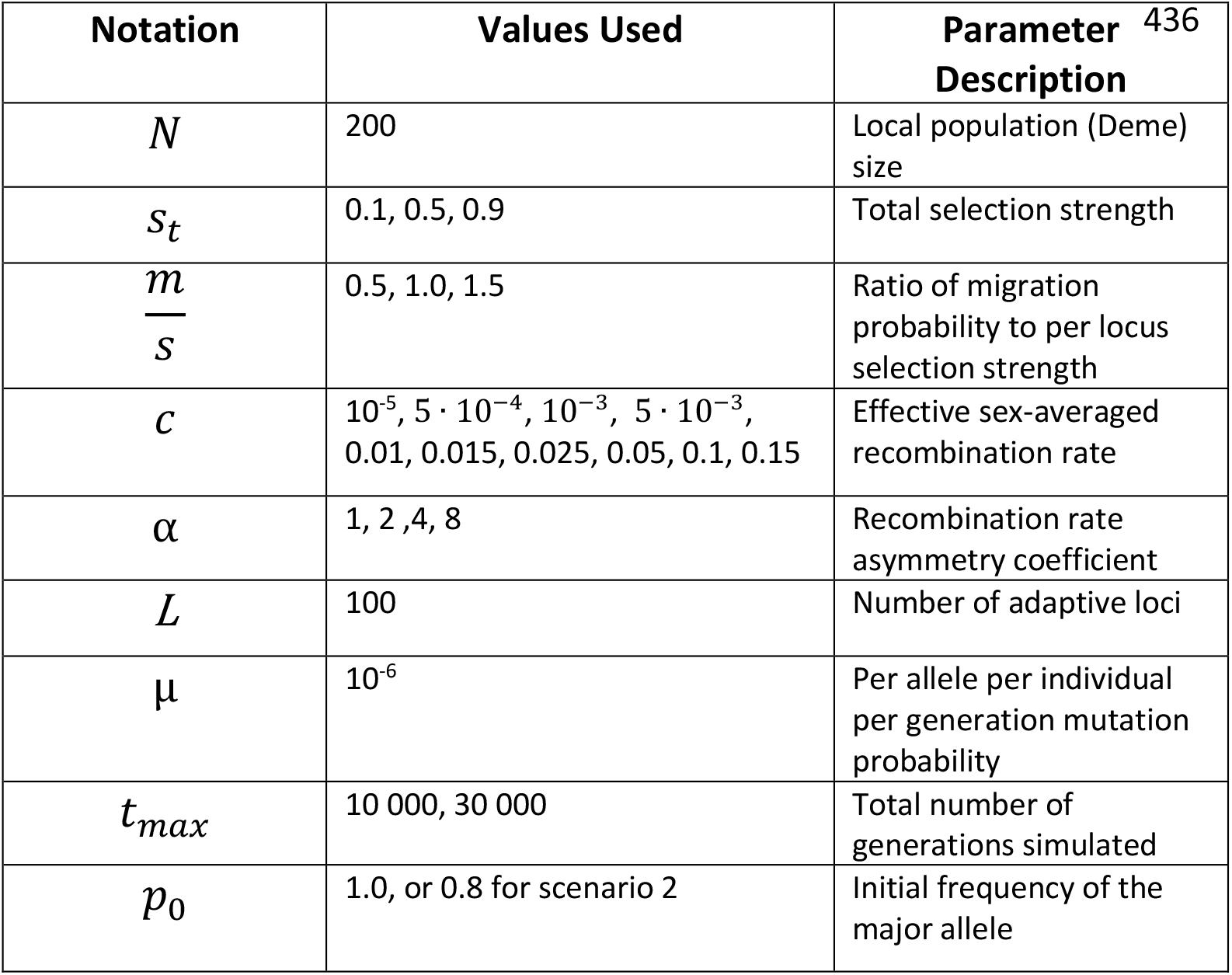
Parameter notations, their explanations, and the values used in the main scenarios.

| Notation | Values Used | Parameter Description <sup>436</sup> |
| --- | --- | --- |
| $N$ | 200 | Local population (Deme) size |
| $s_t$ | 0.1, 0.5, 0.9 | Total selection strength |
| $\frac{m}{s}$ | 0.5, 1.0, 1.5 | Ratio of migration probability to per locus selection strength |
| $c$ | $10^{-5}$, $5 \cdot 10^{-4}$, $10^{-3}$, $5 \cdot 10^{-3}$, 0.01, 0.015, 0.025, 0.05, 0.1, 0.15 | Effective sex-averaged recombination rate |
| $\alpha$ | 1, 2, 4, 8 | Recombination rate asymmetry coefficient |
| $L$ | 100 | Number of adaptive loci |
| $\mu$ | $10^{-6}$ | Per allele per individual per generation mutation probability |
| $t_{max}$ | 10 000, 30 000 | Total number of generations simulated |
| $p_0$ | 1.0, or 0.8 for scenario 2 | Initial frequency of the major allele |

Selection in the two demes is assumed to be divergent, such that at each locus, allele *A* is beneficial in deme 1 and deleterious in deme 2, whereas allele *a* is deleterious in deme 1 and beneficial in deme 2.

Each locus is assumed to contribute to the fitness depending on the individual’s genotype at that locus. For individuals in deme 1, the contributions are 1 + *s*, 1, and 1 − *s*, for the homozygote *AA*, the heterozygote *Aa*, and the homozygote *aa*, respectively. In deme 2, the contributions are instead 1 + *s*, 1, and 1 − *s*, for the homozygote *aa*, the heterozygote *Aa*, and the homozygote *AA*, respectively. For simplicity, the parameter *s* is assumed to be the same for all loci.

We further assume that single-locus genotypes contribute multiplicatively to the individuals’ fitness. Thus, the total fitness of an individual with all locally beneficial, or with all locally deleterious alleles at all *L* loci is given by *w*_*max*_= (1 + *s*)^*L*^, or *w*_*min*_ = (1 − *s*)^*L*^, respectively. Thus, an individual that is perfectly adapted to one deme will suffer a fitness cost (hereafter referred to as total selection, *s*_*t*_) in the opposite deme relative to the individuals that are locally perfectly adapted, amounting to 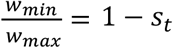. From this, it follows that the per-allele selection coefficient, *s*, can be expressed in terms of the total selection coefficient, *s*_*t*_, as 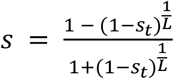 In our simulations we used different values of *s*_*t*_ to account for relatively weak (*s*_*t*_ = 0.1), intermediate (*s*_*t*_ = 0.5), and strong divergent selection (*s*_*t*_ = 0.9; corresponding to *w*_*max*_= 10_*min*_;

Table 1). After the fitness of all individuals has been calculated, *N* female and *N* male gametes are produced as follows. The probability that an individual of a given sex in a given deme produces offspring is given by the fitness of that individual relative to the total fitness of all individuals of the same sex in that deme. Using these probabilities, we sample randomly (with replacement) a total of *N* mothers and *N* fathers (contributing one gamete each) among all available females and males in each deme.

After selection, we perform recombination. The probability of recombination between any consecutive pairs of loci is based on the effective sex-averaged recombination rate parameter, *c*, and the recombination rate asymmetry parameter, *α*. In each simulation both *c* and *α* are assumed to be constant over time. Denoting by *c*_*m*_, and *c*_*f*_ = *αc*_*m*_ the probability of recombination between any consecutive pair of loci in males, and females, respectively, it follows that the effective sex-averaged recombination rate is 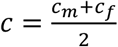.Expressing *c*_*m*_ and *c*_*f*_ in terms of *c* and *α* yields: 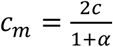, and 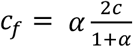. For each parental individual we simulate recombination between loci such that the recombination between each pair of consecutive loci in females (males) occurs independently of each other and with probability *c*_*f*_ (*c*_*m*_).

After recombination, each parent, thus, produces two (recombined) haploid gametes, and we sample uniformly at random one of the two to retain in the gamete pool (both being equally likely). The remaining gamete is discarded.

Mating is assumed to be local and hence it is implemented separately for each deme by randomly pairing, without replacement, the *N* maternal and *N* paternal gametes produced in the previous step. This produces *N* zygotes. After mating, half of the zygotes are randomly assigned the female sex and half are randomly assigned the male sex. Thereafter, we allow for zygote mutation. Mutation is assumed to be symmetric such that the per-generation per-allele probability (*μ*) of allele *A* mutating to allele *a* is equal to the per-generation per-allele probability of allele *a* mutating to allele *A*. In the model, *μ* is chosen in such a way that mutations occur relatively rarely (Table 1).

After this step, all adult individuals die, and zygotes become the new generation of adults. This completes the individuals’ life cycle.

To assess the effect of the initial conditions in our model, we simulated two main different scenarios, as explained next.

### Scenarios

We explored two main scenarios corresponding to primary divergence. In these scenarios, the genomes consisted of *L* = 100 loci and both scenarios used the same parameter values, except for the amount of the genetic variation at the start of the simulations. Namely, we simulated primary divergence without, and with an initial local standing genetic variation (details below).

For comparative purposes with the model in Venu et al. (2024), we additionally simulated a scenario with only three loci under selection. This scenario is further explained in Appendix A. Hereafter, we focus on the former two, main scenarios.

### Main simulated scenarios

In scenario 1 we assume that, for a long period of time, the two demes have been evolving neutrally in a geographic isolation from each other so that alleles at all loci have been driven to fixation within each deme prior to the removal of the geographical barrier, i.e., at the start of the simulations. In this scenario, thus, there is no local standing genetic variation at the start of the simulations. To achieve this, for each individual run, the locally fixed allele (*A* or *a*) at a given locus is chosen uniformly at random (with probability 0.5) and independently for each locus and independently for deme 1 and deme 2. This means that approximately half of the loci simulated are globally monomorphic and the other half are globally polymorphic with the expected allele frequency of 0.5. This also means that within each deme, approximately half of the fixed alleles are beneficial, and half are deleterious. In this scenario, thus, there are no heterozygotes within the demes at the start of the simulations.

In scenario 2 we assume that there is local standing genetic variation at the start of the simulations, and that the initial allele frequencies are equal in both demes. This scenario mimics the arrival of populations to two demes from an old, large (and the same) source population. In this scenario the local frequency of the major allele is (arbitrarily) set to 0.8 at all loci. For each locus and run, the major allele is chosen randomly between the two possible alleles (*A* and *a*). The major allele at a given locus is the same in both demes. The two alleles with such predefined frequencies (0.8, and 0.2 for the major, and minor allele, respectively) are then distributed randomly among the individuals in each deme.

For both scenarios, we ran most of the simulations for 10 000 generations after the initialisation, but in some cases we also assessed the simulation outcomes 30 000 generations after the initialisation (Appendix B). For each parameter set (Table 1), we ran 100 independent simulations.

### Simulation output statistics

At regular intervals (every 50^th^ generation), the local allele frequencies are recorded (but also the allele establishment probability for scenario 3; Appendix A). To quantify the extent of divergence between the two local populations, we computed for each locus the per-generation between-demes allele frequency difference of the allele that is beneficial in deme 1 and deleterious in deme 2 (i.e., allele A) and averaged this quantity over all loci. This quantity is recorded for each run separately. In our results, we present the 5^th^ and 95^th^ percentiles of this quantity, together with its average value over all replicate runs for each parameter set.

## Results

In what follows, we present the results from our two main scenarios with polygenic adaptation. The results from the third scenario (most like that in Venu et al. (2024)) are discussed in Supplementary Material.

### Divergence Without Standing Genetic Variation

As expected (Bulmer, 1972; Yeaman & Otto, 2011; Yeaman & Whitlock, 2011; Rafajlović et al., 2016) our results show that stronger migration hinders divergence, whereas stronger selection facilitates it (compare in Figs. 1-2 and Figs. S1-S2 the speed of divergence, and the extent of the divergence attained at the end of the simulations for different values of *m*, and *s*). The role of the effective recombination rate is more complex. Beyond a threshold recombination rate (which depends on the other model parameters), the extent of divergence initially increases, reaching a maximum, and then starts decreasing (Figs. 1-2, S1-S2). When the recombination rate increases beyond this threshold, the maximum divergence attained becomes smaller, and the speed of loss of divergence is faster (compare Fig.1Q and Fig.1R). However, when the recombination rate is below the threshold value, high divergence is either maintained in a quasi-stationary state by the end of the simulations, or it shows an increasing trend (Fig.1). In this parameter space, divergence is slower when the recombination rate is smaller.

**Figure 1.**
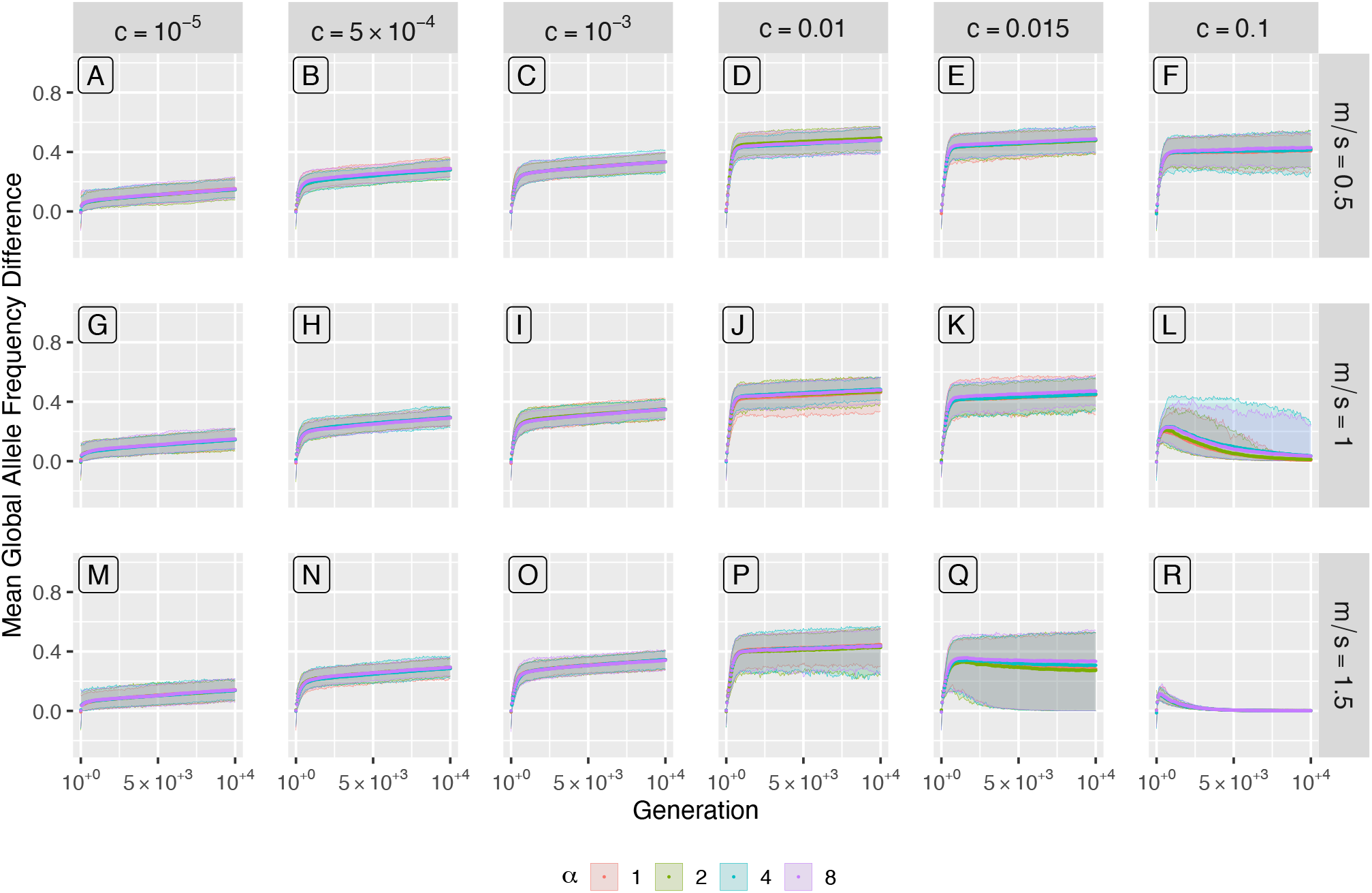
Results under the model of primary divergence **without** local standing genetic variation (scenario 1) under strong selection, s_t_ = 0.9. Shown is the mean global allele frequency difference as a function of time, over 10 000 generations. Panels show results for different values of the effective recombination rate c (increasing left to right) and ratio of migration probability to selection strength m/s (increasing top to bottom). Colours correspond to different values of the recombination rate asymmetry coefficient α: α = 1 (red), α =2 (green), α = 4 (cyan), α = 8 (purple). Ribbons show 5^th^ to 95^th^ percentiles. Results for the additional parameter values used in our simulations are shown in Fig.2 and Figs.S1, S2.

**Figure 2.**
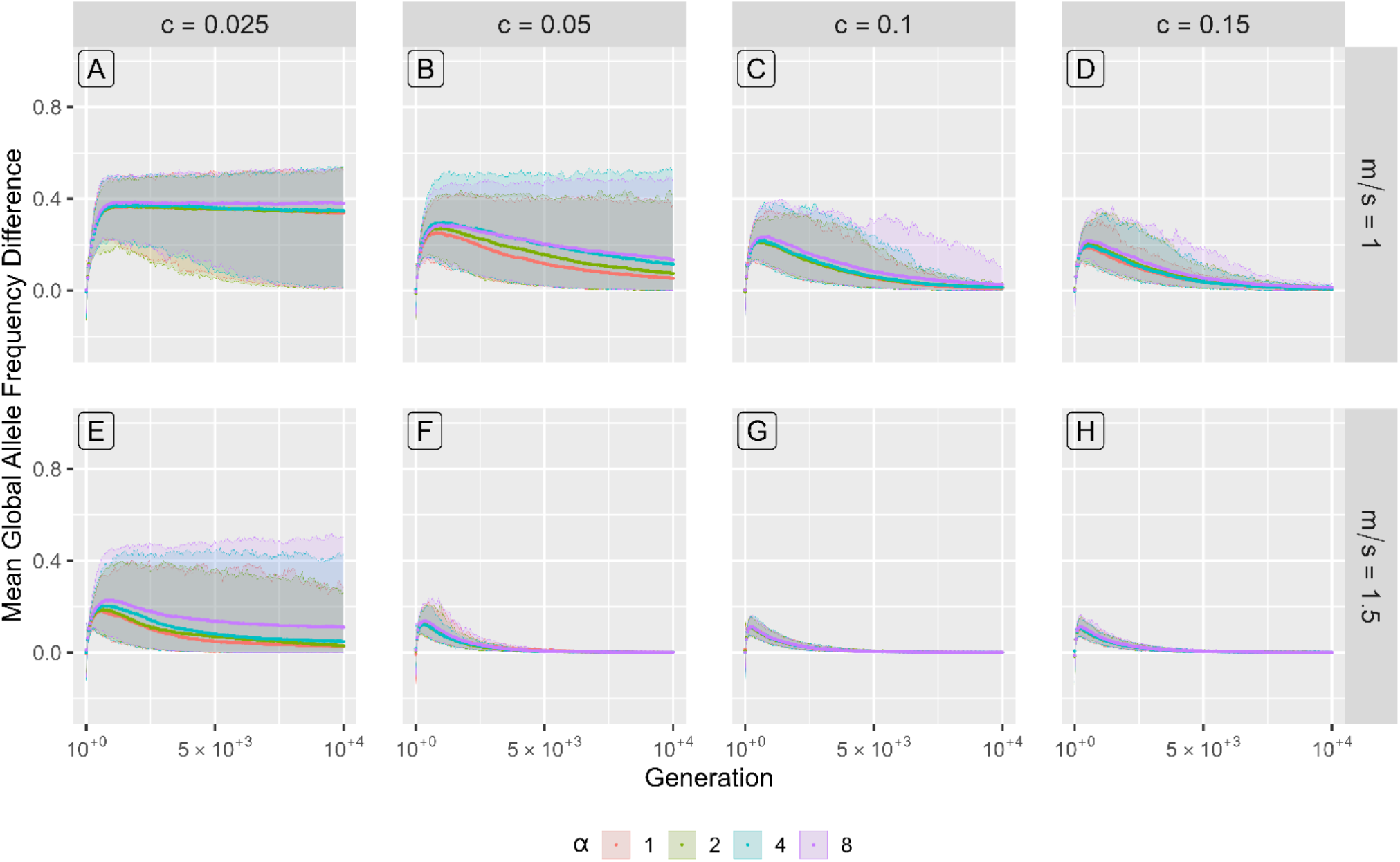
Additional results under the model of primary divergence **without** local standing genetic variation (scenario 1) under strong selection, s_t_ = 0.9. Shown is the mean global allele frequency difference as a function of time, over 10 000 generations. Panels show results for different values of the effective recombination rate c (increasing left to right) and ratio of migration probability to selection strength m/s (increasing top to bottom). Colours correspond to different values of the recombination rate asymmetry coefficient α: α = 1 (red), α =2 (green), α = 4 (cyan), α = 8 (purple). Ribbons show 5^th^ to 95^th^ percentiles. This figure corresponds to Fig. 1, but for additional parameter values. The results for the remaining parameter values are shown in Fig.1 and Fig.S1.

These results are in agreement with the finding of optimal recombination rate by Aeschbacher & Bürger (2014) and the negative effects of strong recombination and drift during divergence (Rafajlović et al. 2016).

The strength of the recombination rate asymmetry between the sexes (α) generally had no discernible effect on divergence (Figs.1-2), except in six combinations of parameter values, all of which were under strong selection (*s*_*t*_ = 0.9) and moderate to strong migration 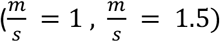. Under moderate migration, 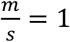, the values of *c* that produced a visible effect were 0.05, and 0.1 (Figs.1L, 2B, 2C), although two additional values of *c* (0.025 and 0.15) also produced some effect, *albeit* weaker (Fig.2A, D). Under strong migration, 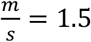, α had an effect on divergence for *c* = 0.015 (Fig. 1Q) and *c* = 0.025 (Fig. 2E). In all cases where α had an effect on divergence, we found that *α* ≥ 4 consistently resulted in the highest extent of divergence at the end of the simulations (Fig. 1L, 1Q, Fig.2A, 2B, 2C, 2E). Interestingly, in only one of these cases (i.e., for s_*t*_ = 0.9, 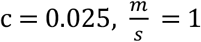), relatively high divergence was retained by the end of the simulated period and it appeared to be in an approximate quasi-stationary state (or with a slow increasing trend for *α* = 8) (Fig.2A and Fig.S3), whereas in the remaining cases, divergence was very low or non-existent by the end of the simulations (Figs. 1-2 and Figs. S1-S2). A similar pattern was seen when the simulations were run for 30 000 generations (Figs. S3, S4).

### Divergence With Standing Genetic Variation

The qualitative effects of migration, selection and the effective recombination rate observed in scenario 1 were retained in scenario 2, but more parameter values resulted in high divergence compared to scenario 1 (compare Fig.3 vs. Fig.1 and Fig.2 vs. Fig.4). Moreover, the recombination rate asymmetry parameter (*α*) had an effect on divergence when both selection and migration were strong (i.e., *s*_*t*_ = 0.9, and 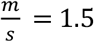), and the effective recombination rate was comparatively high (i.e., c = 0.05, 0.1, 0.15; Figs. 3R, 4F, 4G, 4H and Figs. S5I, S6B). Only in one of these cases (c = 0.05), divergence was retained at a high and approximately quasi-stationary value by the end of the simulations (Fig. 4F), whereas it showed a decreasing trend in the remaining cases. As in scenario 1, these trends were even clearer when the simulations were allowed to run for 30 000 generations (Figs. S7, S8).

**Figure 3.**
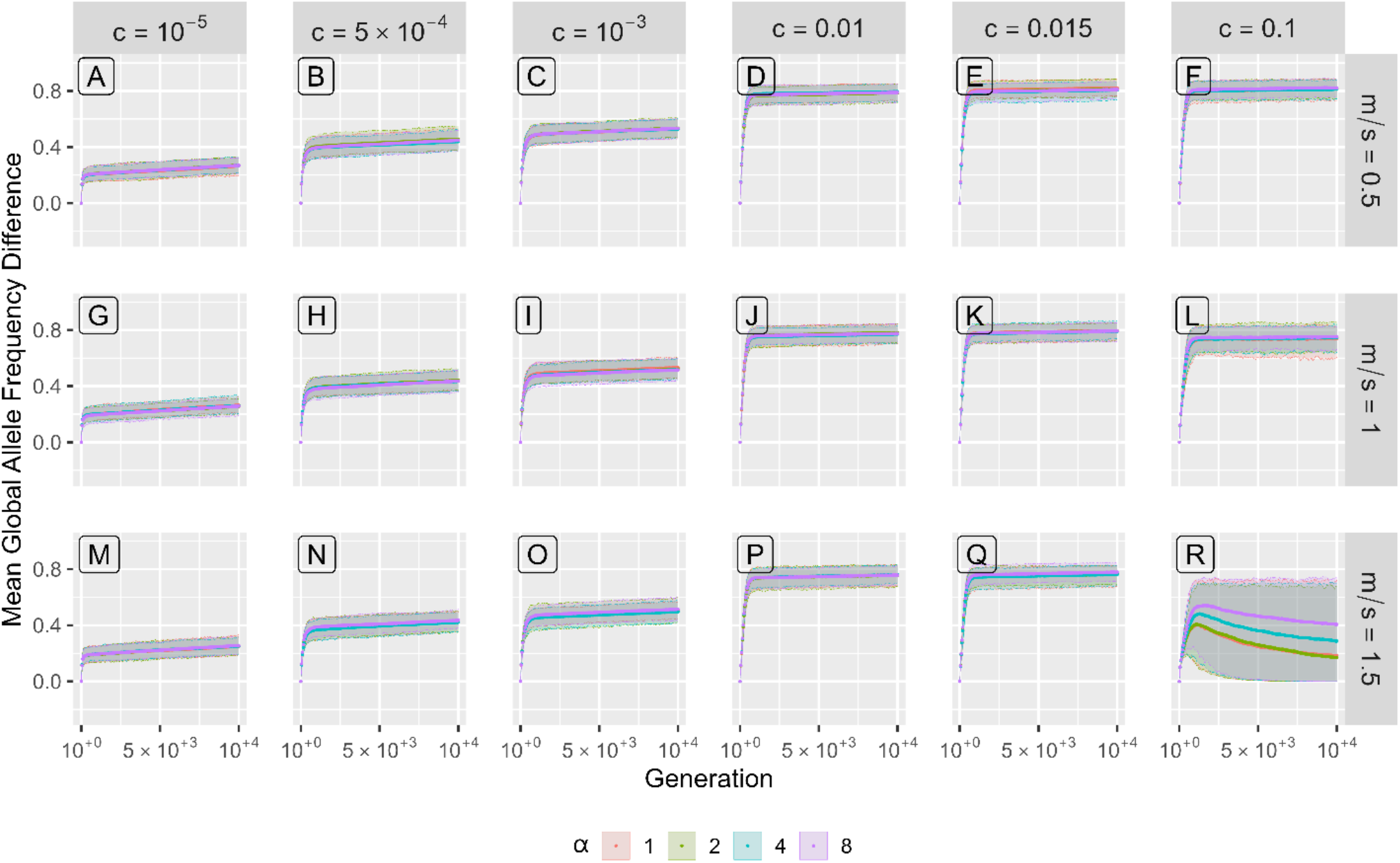
Results under the model of primary divergence **with** local standing genetic variation (scenario 2) under strong selection, s_t_ = 0.9. Shown is the mean global allele frequency difference as a function of time, over 10 000 generations. Panels show results for different values of the effective recombination rate c (increasing left to right) and ratio of migration probability to selection strength m/s (increasing top to bottom). Colours correspond to different values of the recombination rate asymmetry coefficient α: α = 1 (red), α =2 (green), α = 4 (cyan), α = 8 (purple). Ribbons show 5^th^ to 95^th^ percentiles. Results for the additional parameter values used in our simulations are shown in Fig.4 and Figs.S5, S6, S7.

**Figure 4.**
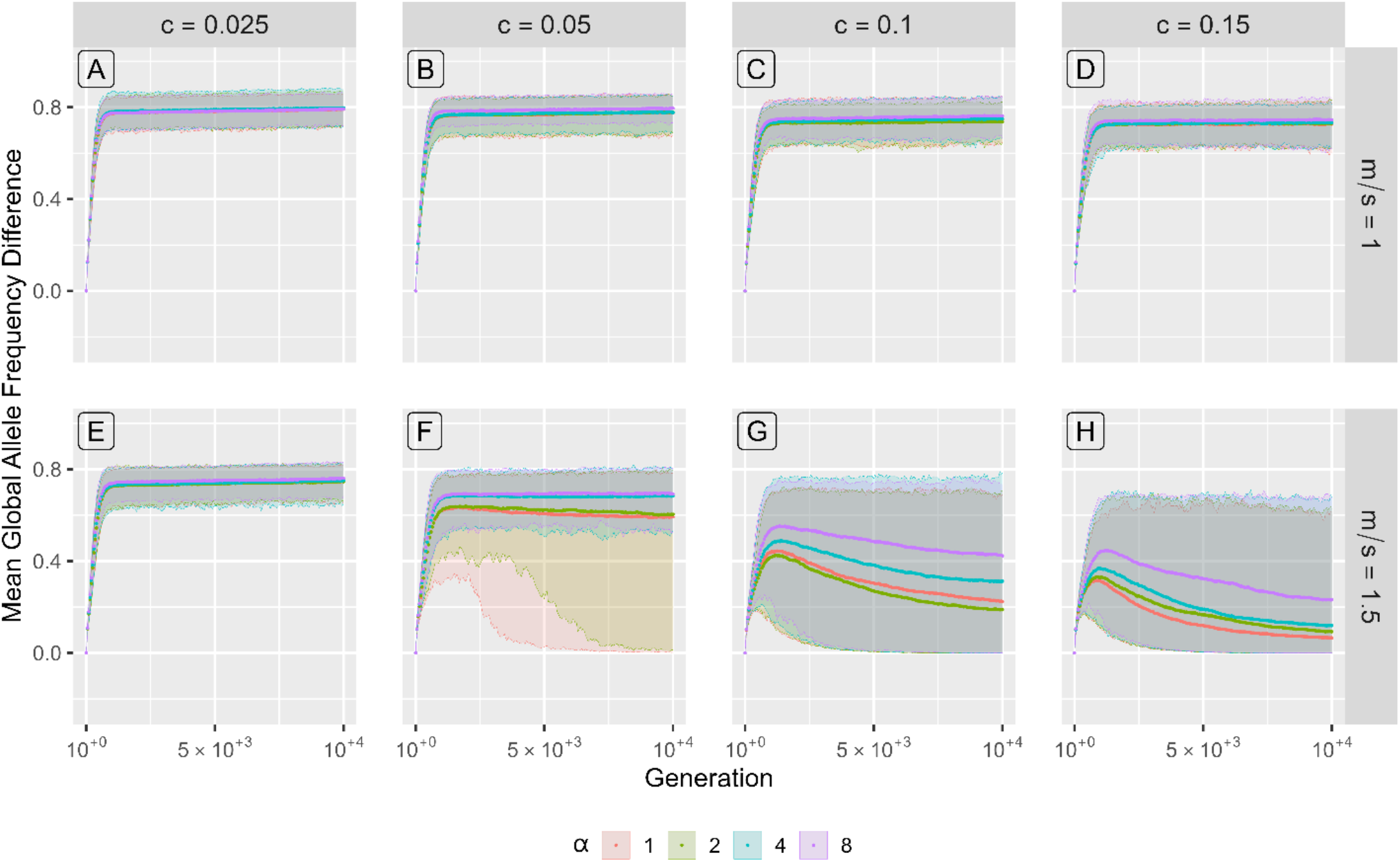
Additional results under the model of primary divergence **with** local standing genetic variation (scenario 2) under strong selection, s_t_ = 0.9. Shown is the mean global allele frequency difference as a function of time over, 10 000 generations. Panels show results for different values of the effective recombination rate c (increasing left to right) and ratio of migration probability to selection strength m/s (increasing top to bottom). Colours correspond to different values of the recombination rate asymmetry coefficient α: α = 1 (red), α =2 (green), α = 4 (cyan), α = 8 (purple). Ribbons show 5^th^ to 95^th^ percentiles. This figure corresponds to Fig. 3, but for additional parameter values.

## Discussion

The role of recombination is dual in the sense that strong recombination can facilitate adaptation by allowing for purging of deleterious alleles, but it can also act to break up locally beneficial allele complexes. On the other hand, weak recombination (i.e., tight linkage) can delay or preclude purging of deleterious alleles, but it can also effectively shield locally beneficial allele complexes from breaking up. The intricate interplay between weak and strong recombination along the genome and within populations can have important consequences on populations’ adaptive dynamics, but they are yet to be comprehensively understood.

It is well known that the rate of recombination varies between individuals and within genomes (Lenormand & Dutheil, 2005; Kong et al., 2010; Martin et al., 2019; Barboza & Blackmon, 2025). One layer of such between-individual variation is seen in sex chromosomes where recombination occurs normally between sex chromosomes of the homogametic sex (XX or ZZ) and no (meaningful) recombination occurs between sex chromosomes of the heterogametic sex (XY or ZW).

Another layer involves sex differences in recombination rates on a fraction of autosomal chromosomes, with more frequent recombination observed in females than in males, e.g., in humans (Bhérer et al., 2017), sticklebacks (Venu et al. 2024), mice and rats, (Dunn, 1920), *Drosophila* (McKee et al., 2012), and many others (Dunn & Bennett, 1967; Lenormand & Dutheil, 2005). In many species, such as mice, different parts of the genome have opposite sex-biased recombination rates although the overall rate remains higher in females (Dunn & Bennett, 1967). For *Bombyx (Shi et al*., *1995)*, a genus of silk moths where females are the heterogametic sex, and some plants (that lack sex chromosomes altogether) (Lenormand & Dutheil, 2005), the situation is reversed, with higher overall rates of recombination in male meiosis. *Bombyx*, and *Drosophila* present an extreme case, *achiasmy*, where recombination is completely absent in the heterogametic sex (i.e., in females, and males, respectively; Lenormand & Dutheil, 2005). However, the heterogametic sex is not always the sex with lower or absent recombination (Dunn & Bennett, 1967).

Notably, sex chromosomes are inherently different from autosomes, and so one may expect that the evolutionary implications of sex differences in recombination rate on autosomes and sex chromosomes can be different. Unlike autosomes, sex chromosomes X and Y (or Z and W) are not homologous. Their size and gene content are markedly different, with the chromosome unique to the heterogametic sex (Y or W) often containing very few genes that are typically not essential, except for sex-determining genes (Charlesworth & Charlesworth, 2025). Furthermore, recombination is functionally absent between X and Y (or Z and W) (Lenormand & Dutheil, 2005) and often highly reduced between homologous X (or Z) chromosomes (Fraïsse & Sachdeva, 2021). In addition, the strength and direction of selection on genes on the X (or Z) chromosome can be sex dependent with some alleles potentially beneficial in females and deleterious in males, and vice versa (Haldane, 1922; Hedrick, 2007; Connallon et al., 2018; Fraïsse & Sachdeva, 2021).

Although a large body of theory has been devoted to the role of differences between the sex chromosomes on adaptation and divergence (Haldane, 1922; Charlesworth et al., 1987; Fraïsse & Sachdeva, 2021, and references therein), much less is known about a potential role of sex differences in autosomal recombination rate. This is an important knowledge gap because many adaptive loci in most species are harboured by autosomes (Lasne et al. 2017).

The aim of this study was to theoretically assess if sex differences in autosomal recombination rate, in the absence of any other sex asymmetries, can facilitate divergence. To achieve this, we used stochastic individual-based simulations of polygenic adaptation and divergence with migration. We assessed model results in dependence of the relative recombination rate between females and males (i.e., the recombination rate asymmetry parameter, *α*) for a range of values of selection strength, migration rate and effective (sex-averaged) recombination rate. In addition, we assessed model results without and with initial standing genetic variation.

Our results show that sex-differences in autosomal recombination rate may facilitate divergence, as argued by Venu et al. (2024). However, our findings indicate that this is an exception rather than a rule. Indeed, the effect is typically weak and only observable under very specific conditions, such as strong asymmetry in recombination rate (i.e., recombination in one sex is at least four times more frequent than in the other sex, *α* ≥ 4), strong selection (*s*_*t*_*=0*.*9*), high effective recombination rate (*c* ≥ 0.025), and intermediate to strong migration, 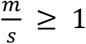. Under these conditions, divergence was, on average, higher and maintained for longer when *α* was larger (Figs. 1L, 1Q, 2B, 2C, 2E, 3R, 4F, 4G, 4H; see also Fig. S1F for *c* = 0.1, Fig. S2A for *c* = 0.05, Fig. S2B for *c* = 0.025, Fig. S3A, S3B, S3C, S3E, S4A for 0.025 ≤ *c* ≤ 0.05, Fig. S4B for *c* = 0.025, Fig. S5I for *c* = 0.1, Fig. S6B for 0.05 ≤ *c* ≤ 0.15, Fig. S7F-H, Fig. S8B for 0.05 ≤ *c* ≤ 0.15). In addition, in some cases without sex differences in recombination rate, divergence was lost within 10000 generations after the start of the simulations, but when *α* was increased above two, loss of divergence was precluded (see Fig. 4F, Fig. S6B for *c* = 0.05). Moreover, with larger values of *α*, cases existed where divergence reached high levels and had an increasing trend by the end of 30 000 generations, despite being lost in cases with *α* ≤ 2 (Figs. S3B, S3E, see also Fig. S4A for *c* = 0.05, and Fig. S4B for *c* = 0.025).

We further found that the effect of recombination rate differences between the sexes tended to increase with increasing *α* and was strongest at the largest value of *α* we used (*α* = 8) although in two of the cases where *α* had an effect on divergence (Figs. 2B, 4F) the effect saturated at *α* = 4.

The results were qualitatively similar in our simulations without and with standing genetic variation, wherein standing genetic variation generally acted to facilitate the divergence and to maintain relatively high divergence levels (compare Figs. 2, 4).

Notably, under the model variant most like that used by Venu et al. (2024) (see Appendix A, and Appendix C) but with only three adaptive loci, two demes, no recombination suppression in heterozygotes, and no asymmetries in migration rates between the sexes (as was assumed in their model), we did not find a support for the positive effect of sex differences in the recombination rate on divergence (Figs. S9-S10). This suggests that the effect of sex differences in recombination rate alone (in the absence of any other processes) is typically very weak (if at all present) except under specific conditions when adaptation is highly polygenic. This is because under a polygenic adaptation a larger number of adaptive loci provides more combinatorial opportunities (Marques et al., 2019) for the dual role of recombination, realised within the separate sexes, to bring about and preserve locally beneficial allele complexes. Such locally beneficial allele complexes are expected to facilitate divergence by reducing gene flow between the populations due to both direct and indirect selection exerted by linkage disequilibrium between the adaptive loci (Barton & Bengtsson, 1986).

In this study, we modelled an extreme case of sex differences in recombination rate, with all males (females) having the same recombination rate. In nature, however, males (females) are likely to follow a distribution of recombination rates. By incorporating this in our model, the recombination rate asymmetry parameter (*α*) would follow a distribution of values (instead of having a single, fixed value, as we assumed here). Based on our results, however, we assert that such a model extension would produce a weaker effect on divergence than that observed in the present study because the overall effect would be smeared out, especially by male-female mates with small differences in recombination rate.

In addition, our model assumed random mating. It is possible that assortative mating between the alike mates would amplify the effect of sex differences in recombination rate, but this remains for future work.

We conclude based on our findings that sex-differences in autosomal recombination rate are, in most cases, unlikely to noticeably contribute to the process of adaptation and divergence. More detailed theoretical studies on the role and evolution of different layers of recombination-rate variation between and within individuals are needed to better understand how they shape local adaptation, and population divergence.

## Data and code availability statement

All computer codes used in this study will be uploaded to Dryad and made publicly available upon the acceptance of the manuscript.

## Author contributions

MR conceived and designed the study. AM wrote the computer code with guidance and input from MR. AM ran the simulations. AM and MR analysed and interpreted the results. AM and MR wrote the first draft of the manuscript.

## Acknowledgments

We should like to acknowledge the financial support by the Swedish Research Council Vetenskapsrådet (grant number 2021-05243) awarded to MR. In addition, we should like to express the gratitude to the Littorina Research Community for their comments on the preliminary results of the study presented in January, 2026.

## References

Aeschbacher, S., & Bürger, R. (2014). The Effect of Linkage on Establishment and Survival of Locally Beneficial Mutations. Genetics, 197(1), 317–336. 10.1534/genetics.114.163477

Barboza, A., & Blackmon, H. (2025). Drivers of achiasmatic meiosis: sexual antagonism versus heteromorphy-dependent aneuploidy across sex-chromosome divergence. G3 Genes|Genomes|Genetics, 15(11). 10.1093/g3journal/jkaf217

Barton, N., & Bengtsson, B. O. (1986). The barrier to genetic exchange between hybridising populations. Heredity, 57(3), 357–376. 10.1038/hdy.1986.135

Bezanson, J., Edelman, A., Karpinski, S., & Shah, V. B. (2017). Julia: A Fresh Approach to Numerical Computing. SIAM Review, 59(1), 65–98. 10.1137/141000671

Bhérer, C., Campbell, C. L., & Auton, A. (2017). Refined genetic maps reveal sexual dimorphism in human meiotic recombination at multiple scales. Nat Commun, 8, 14994. 10.1038/ncomms14994

Bulmer, M. G. (1972). Multiple Niche Polymorphism. The American Naturalist, 106(948), 254–257. 10.1086/282765

Charlesworth, B., Campos, J. L., & Jackson, B. C. (2018). Faster-X evolution: Theory and evidence from Drosophila. Molecular Ecology, 27(19), 3753–3771. 10.1111/mec.14534

Charlesworth, B., & Charlesworth, D. (2025). HJ Muller and the Relationship Between Sex Chromosome Degeneration and the Evolution of Dosage Compensation. Genome Biology and Evolution, 17(11). 10.1093/gbe/evaf195

Charlesworth, B., Coyne, J. A., & Barton, N. H. (1987). The Relative Rates of Evolution of Sex Chromosomes and Autosomes. The American Naturalist, 130(1), 113–146. 10.1086/284701

Connallon, T., Débarre, F., & Li, X.-Y. (2018). Linking local adaptation with the evolution of sex differences. Philosophical Transactions of the Royal Society B: Biological Sciences, 373(1757). 10.1098/rstb.2017.0414

Dunn, L. C. (1920). LINKAGE IN MICE AND RATS. Genetics, 5(3), 325–343. 10.1093/genetics/5.3.325

Dunn, L. C., & Bennett, D. (1967). Sex differences in recombination of linked genes in animals. Genetical Research, 9(2), 211–220. 10.1017/S0016672300010491

Eriksson, M., & Rafajlović, M. (2021). The Effect of the Recombination Rate between Adaptive Loci on the Capacity of a Population to Expand Its Range. The American Naturalist, 197(5), 526–542. 10.1086/713669

Felsenstein, J. (1981). Skepticism Towards Santa Rosalia, or Why are There so Few Kinds of Animals? Evolution, 35(1), 124–138. 10.2307/2407946

Fraïsse, C., & Sachdeva, H. (2021). The rates of introgression and barriers to genetic exchange between hybridizing species: sex chromosomes vs autosomes. Genetics, 217(2). 10.1093/genetics/iyaa025

Haldane, J. B. S. (1922). Sex ratio and unisexual sterility in hybrid animals. Journal of Genetics, 12(2), 101–109. 10.1007/BF02983075

Hedrick, P. W. (2007). SEX: DIFFERENCES IN MUTATION, RECOMBINATION, SELECTION, GENE FLOW, AND GENETIC DRIFT. Evolution, 61(12), 2750–2771. 10.1111/j.1558-5646.2007.00250.x

Kong, A., Thorleifsson, G., Gudbjartsson, D. F., Masson, G., Sigurdsson, A., Jonasdottir, A., Walters, G. B., Jonasdottir, A., Gylfason, A., Kristinsson, K. T., Gudjonsson, S. A., Frigge, M. L., Helgason, A., Thorsteinsdottir, U., & Stefansson, K. (2010). Fine-scale recombination rate differences between sexes, populations and individuals. Nature, 467(7319), 1099–1103. 10.1038/nature09525

Lasne, C, Sgró, C. M., & Connallon, T. (2017). The relative contributions of the X chromosome and autosomes to local adaptation. Genetics, 3(1), 1285–1304. 10.1534/genetics.116.194670

Lenormand, T., & Dutheil, J. (2005). Recombination difference between sexes: a role for haploid selection. PLoS Biol, 3(3), e63. 10.1371/journal.pbio.0030063

Marques, D. A., Meier, J. I., & Seehausen, O. (2019). A Combinatorial View on Speciation and Adaptive Radiation. Trends in Ecology & Evolution, 34(6), 531–544. 10.1016/j.tree.2019.02.008

Martin, S. H., Davey, J. W., Salazar, C., & Jiggins, C. D. (2019). Recombination rate variation shapes barriers to introgression across butterfly genomes. PLOS Biology, 17(2), e2006288. 10.1371/journal.pbio.2006288

McKee, B. D., Yan, R., & Tsai, J. H. (2012). Meiosis in male Drosophila. Spermatogenesis, 2(3), 167–184. 10.4161/spmg.21800

R Core Team. (2025). R: A Language and Environment for Statistical Computing. In R Foundation for Statistical Computing. https://www.R-project.org/

Rafajlović, M., Emanuelsson, A., Johannesson, K., Butlin, R. K., & Mehlig, B. (2016). A universal mechanism generating clusters of differentiated loci during divergence-with-migration. Evolution, 70(7), 1609–1621. 10.1111/evo.12957

Shi, J., Heckel, D. G., & Goldsmith, M. R. (1995). A genetic linkage map for the domesticated silkworm, Bombyx mori, based on restriction fragment length polymorphisms. Genetical Research, 66(2), 109–126. 10.1017/S0016672300034467

Venu, V., Harjunmaa, E., Dreau, A., Brady, S., Absher, D., Kingsley, D. M., & Jones, F. C. (2024). Fine-scale contemporary recombination variation and its fitness consequences in adaptively diverging stickleback fish. Nature Ecology & Evolution, 8(7), 1337–1352. 10.1038/s41559-024-02434-4

Wickham, H. (2016). ggplot2: Elegant Graphics for Data Analysis. Springer-Verlag New York. https://ggplot2.tidyverse.org

Yeaman, S., & Otto, S. P. (2011). ESTABLISHMENT AND MAINTENANCE OF ADAPTIVE GENETIC DIVERGENCE UNDER MIGRATION, SELECTION, AND DRIFT. Evolution, 65(7), 2123–2129. 10.1111/j.1558-5646.2011.01277.x

Yeaman, S., & Whitlock, M. C. (2011). The genetic architecture of adaptation under migration-selection balance. Evolution, 65(7), 1897–1911. 10.1111/j.1558-5646.2011.01269.x

