## Supplementary Information for "Do sex differences in autosomal recombination rates facilitate divergence?"

### Supplementary Material for "Do sex differences in autosomal recombination rates facilitate divergence?"

This Supplementary material consists of three Appendices. In Appendix A, we present model details for our scenario 3, mimicking the model by Venu et al. (2024). In appendix B, we present supplementary figures for our main two scenarios. In appendix C, we show the figures and results we obtained for scenario 3.

#### Appendix A
#### Model details for Scenario 3

In this scenario we modify our model to align more closely with the one described in Venu et al. (2024) (Table S1). Here, there are only three adaptive loci. The initial frequencies of the locally beneficial alleles in deme 1 are 1 for allele A at locus 1, 0 for allele A at locus 2 and 1 for allele A at locus 3. The frequencies of allele A in deme 2 (here locally deleterious) are 0 for all three loci. From generation 0 to generation 29, locus 2 is assumed to evolve neutrally without affecting fitness ($s = 0$), meaning that the individuals in both demes are perfectly adapted at the initialisation. During these initial 29 generations, fitness is calculated based only on loci 1 and 3. Migration occurs from the first generation and onwards. Then, as in Venu et al., (2024), at generation 30, two individuals in deme 1 mutate, so that they are homozygous for the locally beneficial allele at all loci. From generation 30 and onwards, all three loci are assumed to contribute to the fitness of individuals. Then, in line with Venu et al. (2024), to achieve the same difference in fitness between the most and least fit phenotype as before generation 30, the fitness contribution of each locus is lowered. In other words, to keep $s_t$ the same throughout the simulation, $s$ is lowered after generation 29 to account for the additional locus being under selection. For scenario 3, to compare with Venu et al. (2024), the probability of establishment and the average number of generations until a successful establishment of the introduced allele was calculated. This was done as follows. First, for each replicate run, the generation where the frequency of the beneficial mutation at locus 2 in deme 1 rose above 0.5 was noted, and the average frequency of that allele between that generation and the final generation was calculated. If the average frequency was above or equal to 0.45 (to allow for stochastic fluctuations around 0.5), the establishment in that replicate run was deemed as successful. The value 0.45 was chosen arbitrarily to account for random frequency fluctuations around 0.5 due to drift as well as other processes at play. In cases of successful establishment, the time needed for the successful establishment of the beneficial mutation at locus 2 in deme 1 was set to the generation when the frequency first rose above 0.5. Second, for each unique parameter combination, the mean and $5^{th}$ to $95^{th}$ percentiles of the generations at which the successful establishment occurred was determined. Finally, the establishment probability was calculated as the number of the replicate runs in which the establishment of the beneficial mutation at locus 2 in deme 1 was successful divided by the total number of replicate runs.

| Notation | Values Used | Parameter Description |
|---|---|---|
| $N$ | 200 | Local population (Deme) size |
| $s_t$ | 0.1, 0.5, 0.9 | Total selection strength |
| $c$ | $10^{-5}$,$10^{-4}$, $10^{-3}$, $5.5 \cdot 10^{-3}$, 0.01, 0.055, 0.1, 0.2 | Effective sex-averaged recombination rate |
| $\alpha$ | 1, 5, 10, 20 | Recombination rate asymmetry coefficient |
| L | 3 | Number of adaptive loci |
| $\mu$ | 0 | Per allele per individual per generation mutation probability |
| $t_{max}$ | 500 | Total number of generations simulated |
| $p_0$ | At locus 2, deme 1, t = 30: p0 = 0.01. | Initial frequency of the major allele |
| $m$ | 0.1, 0.2, 0.3, 0.4 | Per-individual migration probability |

*Table S1. Table of parameter values used in scenario 3.*

### Appendix B

#### Supplementary Figures for Scenario 1 and 2

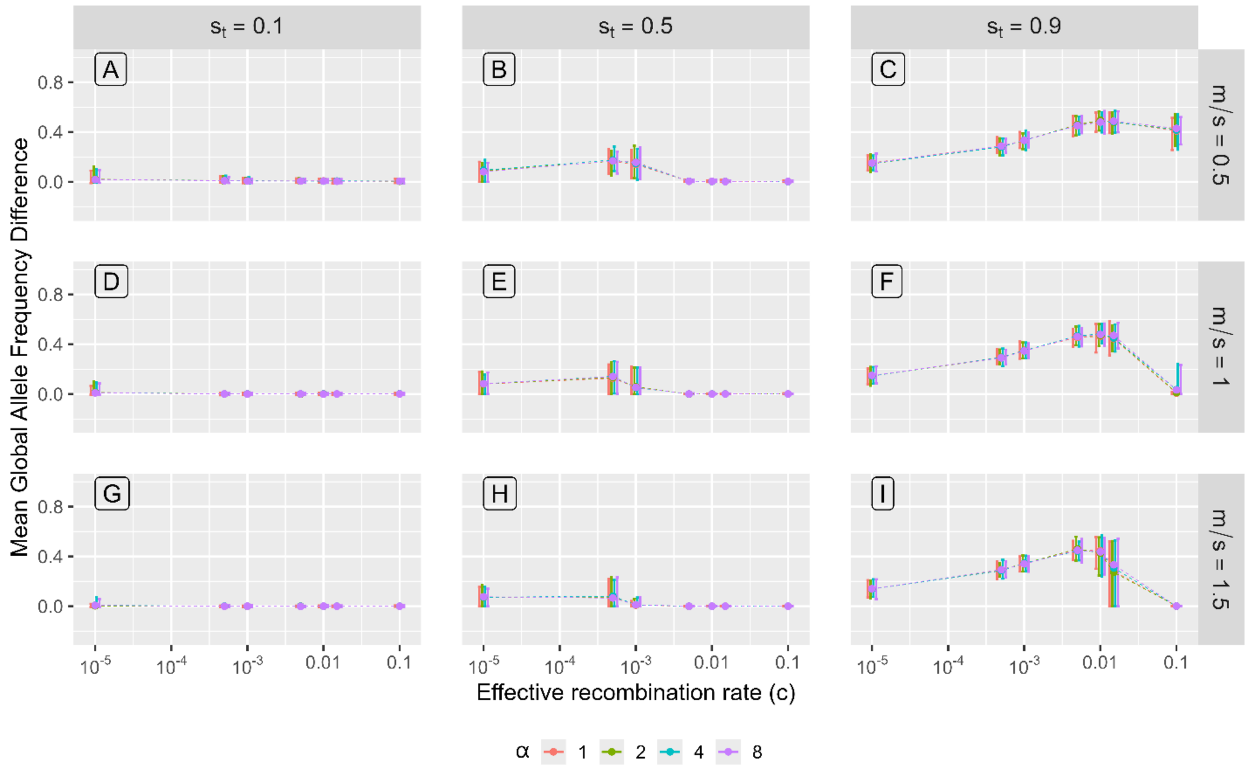


*Figure S1. Results under the model of primary divergence* ***without*** *local standing genetic variation (scenario 1) after 10 000 generations of evolution. Shown is the mean global allele frequency difference attained at generation 10 000 as a function of the effective recombination rate. Results for some values of c are not shown for visibility purposes (note the log10 scale on the x-axis), but they are included in Fig.2 in the main text and Fig.S2 in Appendix B. Panels show the results for different values of the total selection $s_t$ (increasing left to right) and ratio of migration probability to selection strength m/s (increasing top to bottom). Colours correspond to different values of the recombination rate asymmetry coefficient α: α = 1 (red), α =2 (green), α = 4 (cyan), α = 8 (purple). Error bars show 5$^{th}$ to 95$^{th}$ percentiles.*

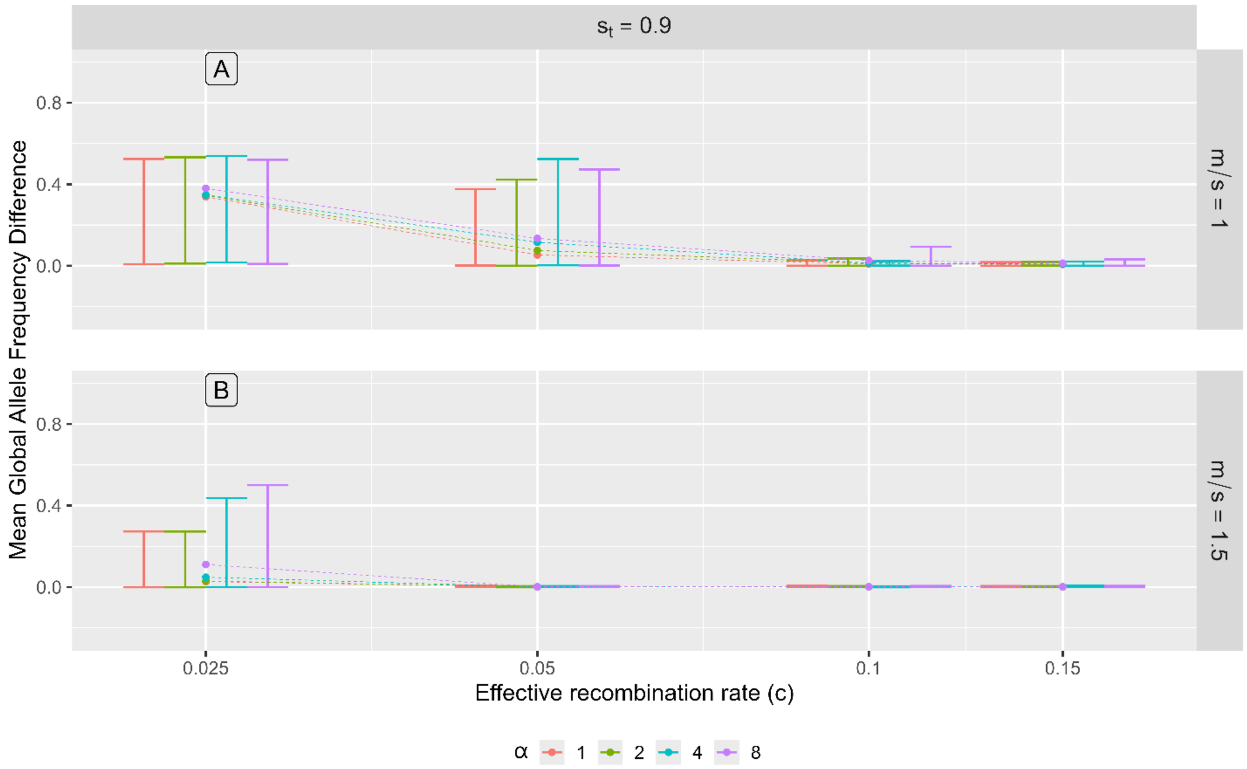


*Figure S2. Results under the model of primary divergence **without** local standing genetic variation (scenario 1) after 10 000 generations of evolution for selected parameter values (results for the remaining parameter values we used are shown in Fig. S1). Shown is the mean global allele frequency difference attained at generation 10 000 as a function of the effective recombination rate. Panels correspond to different values of the ratio of migration probability to selection strength, m/s (increasing top to bottom). Colours correspond to different values of the recombination rate asymmetry coefficient α: α = 1 (red), α =2 (green), α = 4 (cyan), α = 8 (purple). Error bars show $5^{th}$ to $95^{th}$ percentiles.*

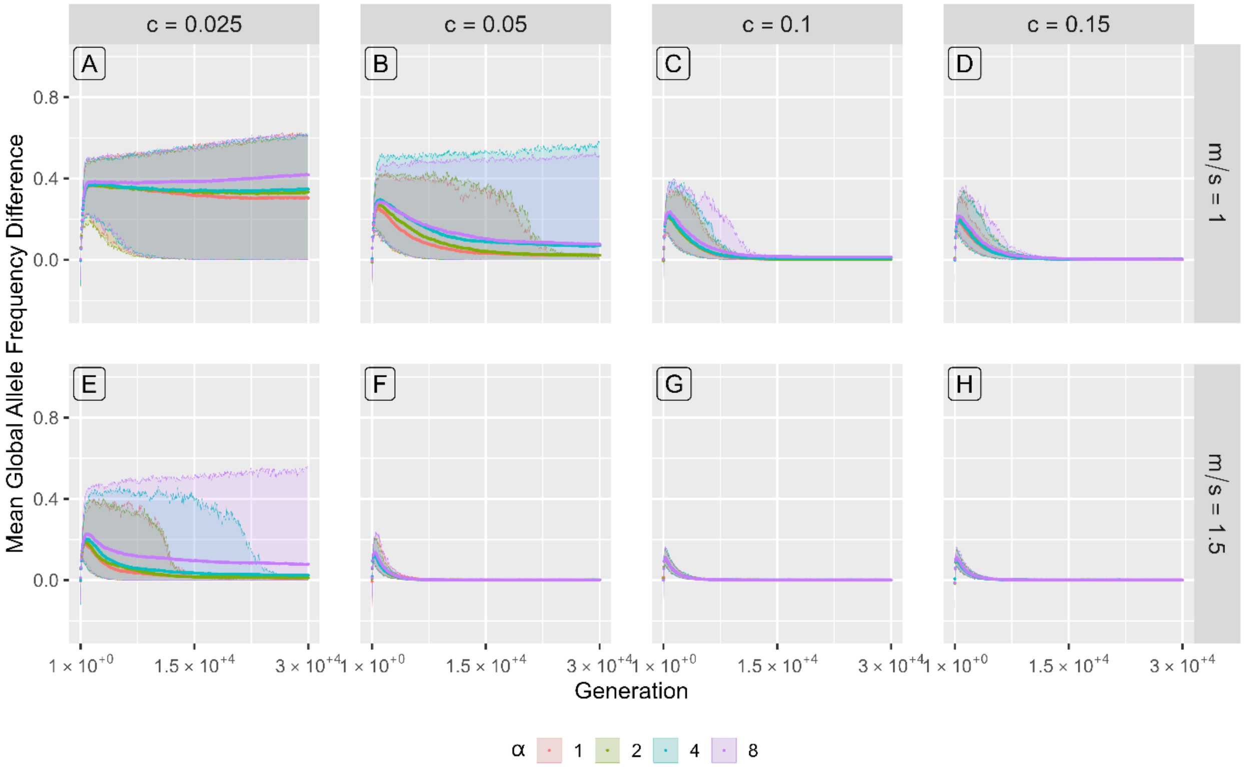


*Figure S3. Additional results under the model of primary divergence* ***without*** *local standing genetic variation (scenario 1) under strong selection, $s_t = 0.9$. Shown is the mean global allele frequency difference as a function of time over 30 000 generations. Panels show results for different values of the effective recombination rate c (increasing left to right) and ratio of migration probability to selection strength m/s (increasing top to bottom). Colours correspond to different values of the recombination rate asymmetry coefficient α: α = 1 (red), α =2 (green), α = 4 (cyan), α = 8 (purple). Ribbons show 5$^{th}$ to 95$^{th}$ percentiles. This figure corresponds to Fig.2, but here the total number of generations is extended to 30 000.*

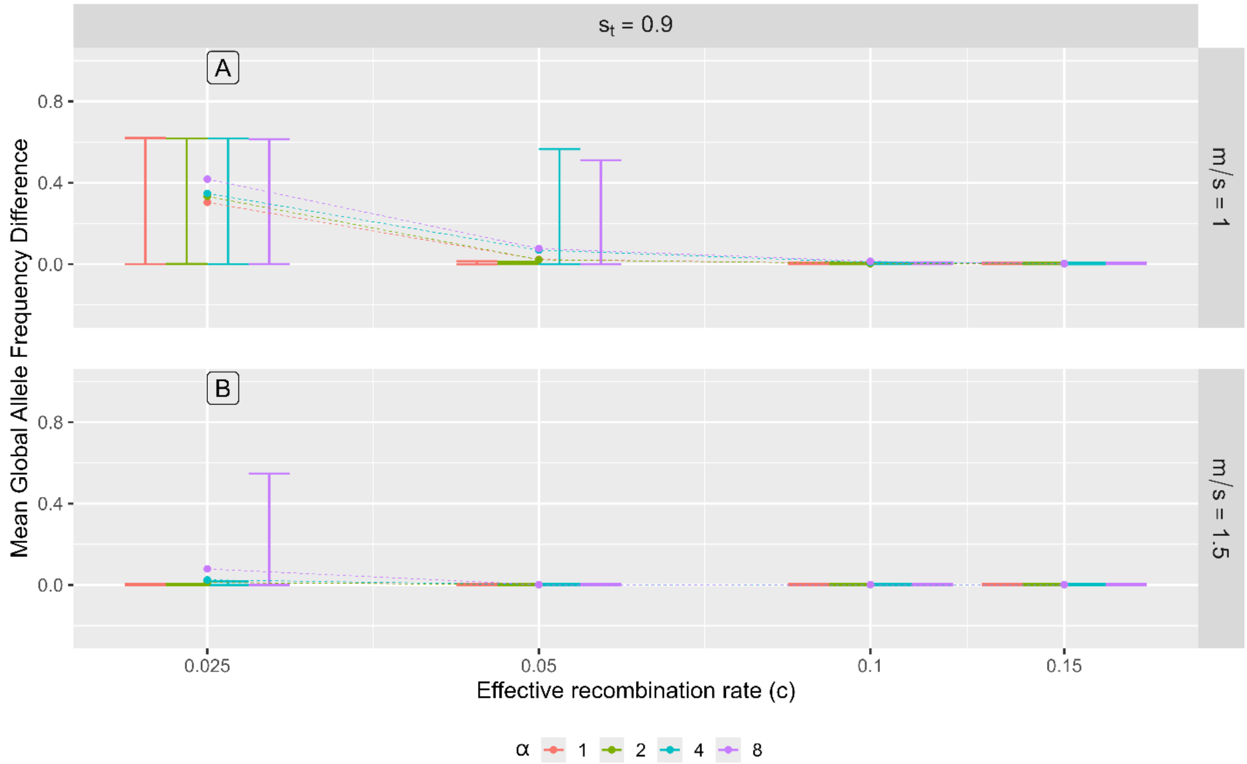


*Figure S4. Results under the model of primary divergence* ***without*** *local standing genetic variation (scenario 1) after 30 000 generations of evolution for selected parameter values (results for the remaining parameter values we used are shown in Fig. S.2). Shown is the mean global allele frequency difference attained at generation 30 000 as a function of the effective recombination rate. Panels correspond to different values of the ratio of migration probability to selection strength, m/s (increasing top to bottom). Colours correspond to different values of the recombination rate asymmetry coefficient α: α = 1 (red), α =2 (green), α = 4 (cyan), α = 8 (purple). Error bars show $5^{th}$ to $95^{th}$ percentiles.*

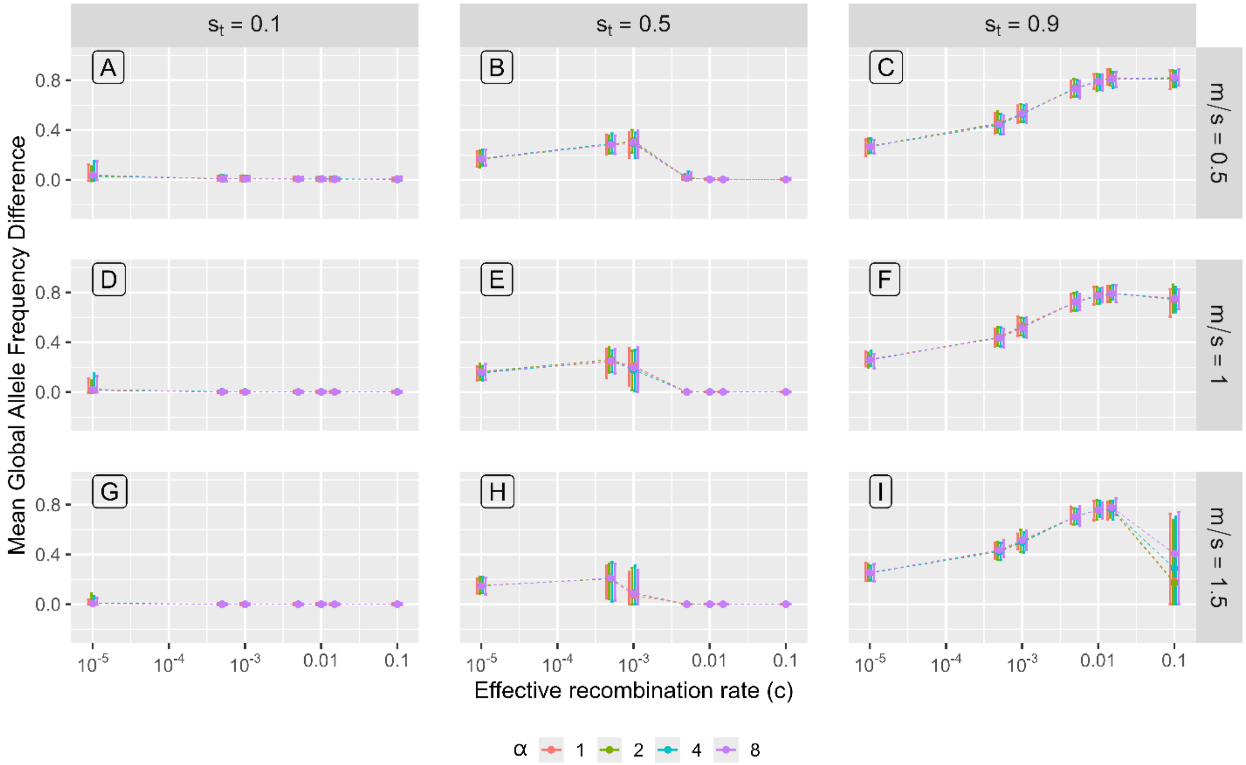


*Figure S5. Results under the model of primary divergence* ***with*** *local standing genetic variation (scenario 2) after 10 000 generations of evolution. Shown is the mean global allele frequency difference attained at generation 10 000 as a function of the effective recombination rate. Results for some values of c are not shown for visibility purposes but they are included in Fig.3 in the main text and Fig.S6 in Appendix B. Panels show the results for different values of the total selection $s_t$ (increasing left to right) and ratio of migration probability to selection strength m/s (increasing top to bottom). Colours correspond to different values of the recombination rate asymmetry coefficient α: α = 1 (red), α =2 (green), α = 4 (cyan), α = 8 (purple). Error bars show 5th to 95th percentiles.*

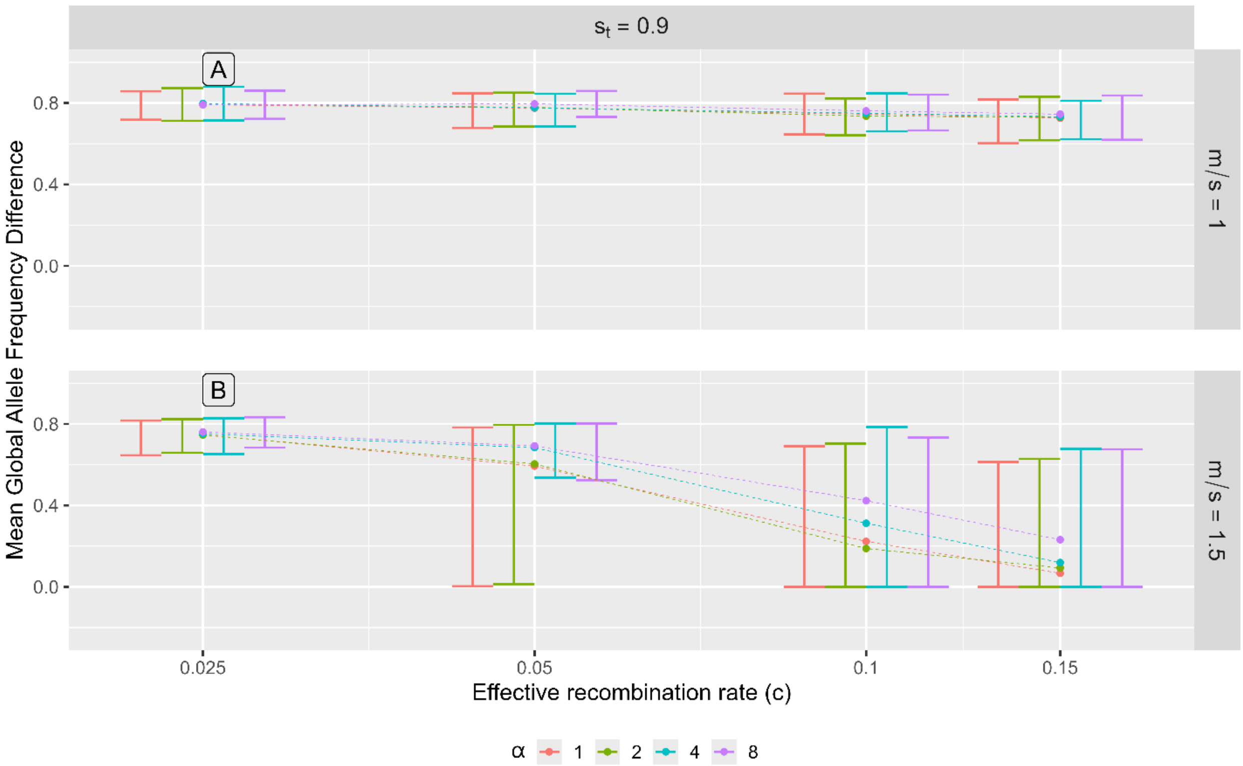


*Figure S6. Results under the model of primary divergence* ***with*** *local standing genetic variation (scenario 2) after 10 000 generations of evolution for selected parameter values (results for the remaining parameter values we used are shown in Fig. S5). Shown is the mean global allele frequency difference attained at generation 10 000 as a function of the effective recombination rate. Panels correspond to different values of the ratio of migration probability to selection strength, m/s (increasing top to bottom). Colours correspond to different values of the recombination rate asymmetry coefficient α: α = 1 (red), α =2 (green), α = 4 (cyan), α = 8 (purple). Error bars show 5^th^ to 95^th^ percentiles.*

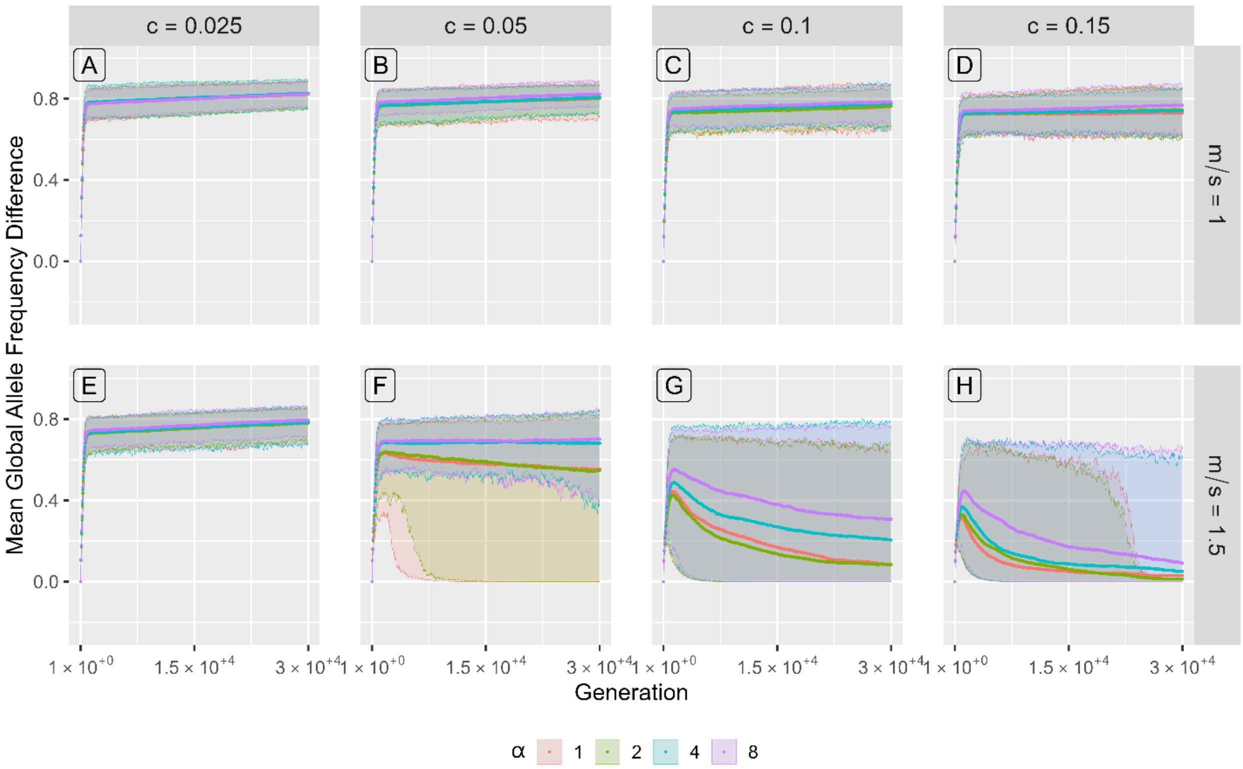


*Figure S7. Additional results under the model of primary divergence **with** local standing genetic variation (scenario 2) under strong selection, $s_t = 0.9$. Shown is the mean global allele frequency difference as a function of time over 30 000 generations. Panels show results for different values of the effective recombination rate c (increasing left to right) and ratio of migration probability to selection strength m/s (increasing top to bottom). Colours correspond to different values of the recombination rate asymmetry coefficient α: α = 1 (red), α =2 (green), α = 4 (cyan), α = 8 (purple). Ribbons show $5^{th}$ to $95^{th}$ percentiles. This figure corresponds to Fig.4, but here the total number of generations is extended to 30 000.*

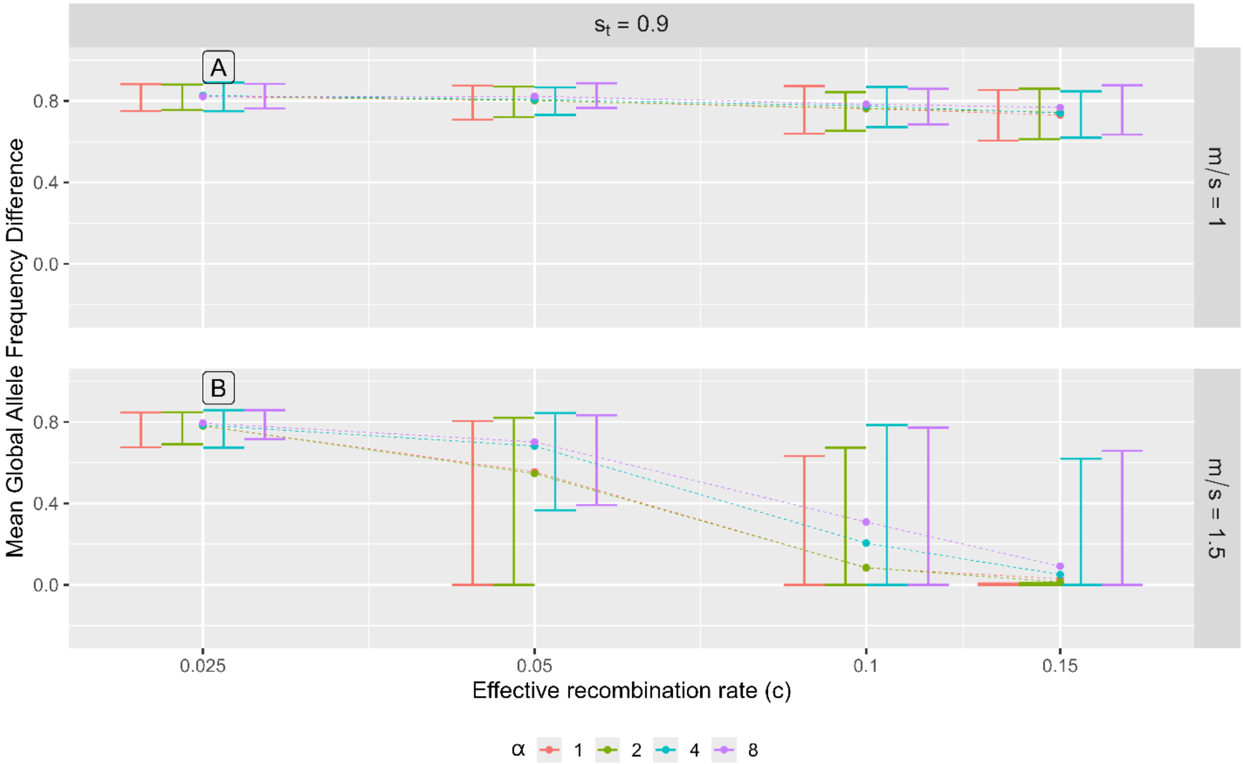


*Figure S8. Results under the model of primary divergence **with** local standing genetic variation after 30 000 generations of evolution (scenario 2) for selected parameter values (results for the remaining parameter values we used are shown in Fig. S5). Shown is the mean global allele frequency difference attained at generation 30 000 as a function of the effective recombination rate. Panels correspond to different values of the ratio of migration probability to selection strength, m/s (increasing top to bottom). Colours correspond to different values of the recombination rate asymmetry coefficient α: α = 1 (red), α =2 (green), α = 4 (cyan), α = 8 (purple). Error bars show $5^{th}$ to $95^{th}$ percentiles. In the case of c = 0.15, α = 1, by 30 000 generations, divergence was lost in most replicate runs except in a few cases, where the divergence attained was relatively high. This resulted in the means being slightly higher than the $5^{th}$ to $95^{th}$ percentiles for this parameter set..*

#### Appendix C

##### Results and Figures for Scenario 3

The establishment probability of the introduced beneficial mutation at locus 2 was positively and strongly associated with the selection strength, weakly negatively associated with the migration rate, and in some cases weakly negatively associated with the effective recombination rate, especially under an intermediate selection and a high effective recombination rate (Figs. S1, S2). Sex-differences in recombination rate produced no visible effect on divergence (Figs. S1, S2).

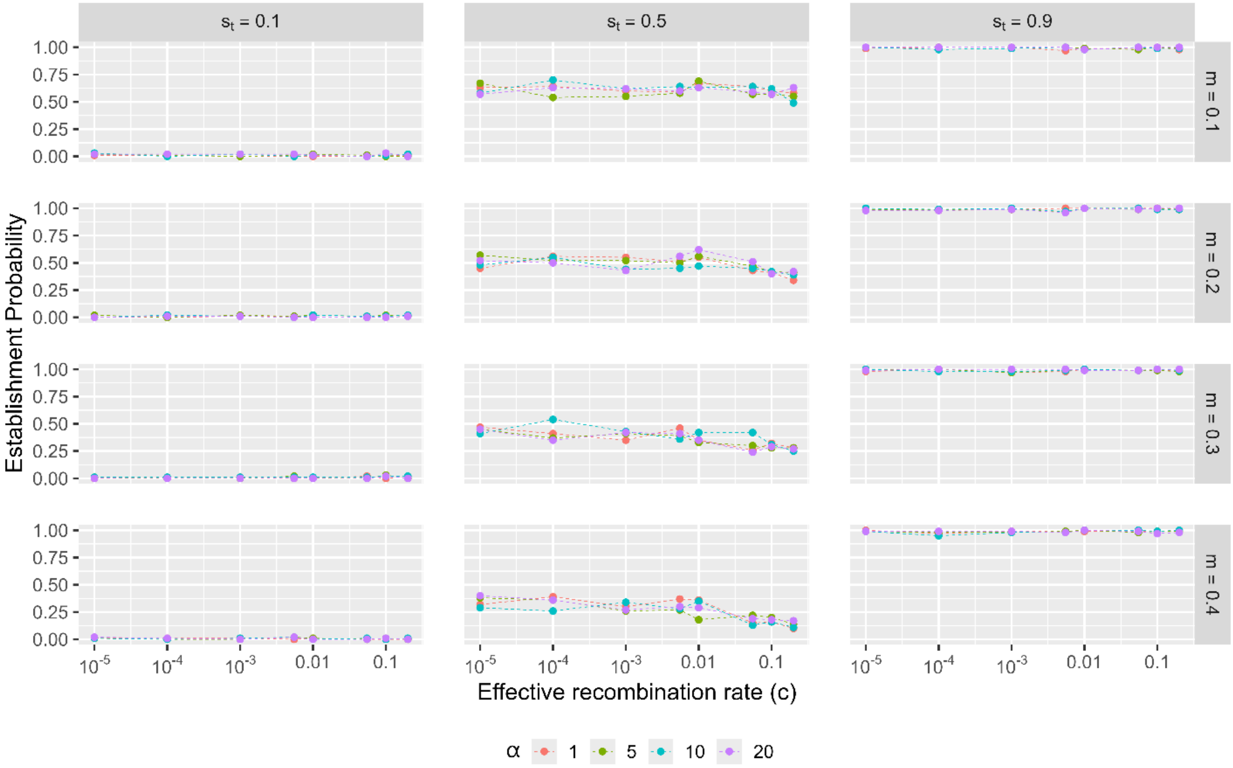


*Figure S9. Results under the model of secondary contact with three loci (scenario 3). Shown is the probability of establishment of the locally beneficial mutation (by generation 500) at locus 2 in deme 1 as a function of the effective recombination rate. Panels show the results for different values of the total selection $s_t$ (increasing left to right) and ratio of migration probability to selection strength m/s (increasing top to bottom). Colours correspond to different values of the recombination rate asymmetry coefficient α: α = 1 (red), α =5 (green), α = 10 (cyan), α = 20 (purple). Error bars show 5$^{th}$ to 95$^{th}$ percentiles.*

.

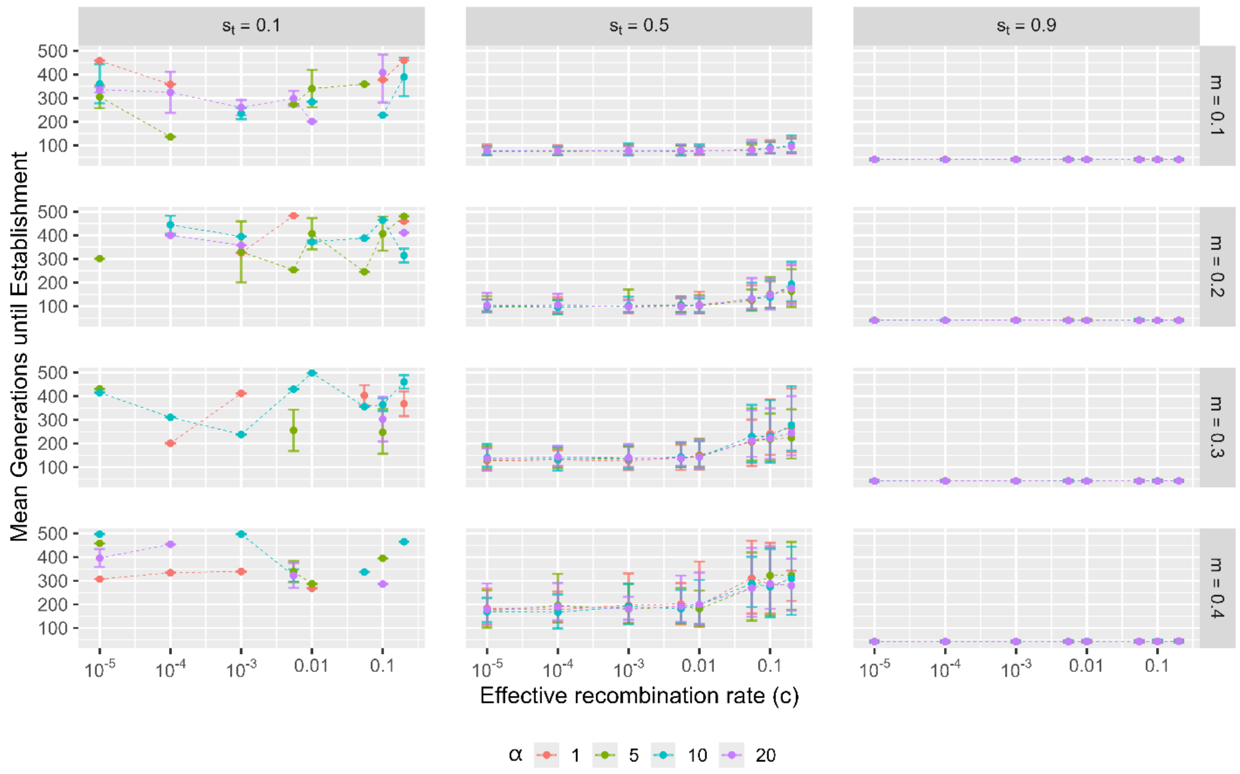


*Figure S10. Results under the model of secondary contact with three loci (scenario 3). Shown is the mean number of generations until establishment (conditioned on cases where establishment was successful) of the locally beneficial mutation at locus 2 in deme 1 as a function of the effective recombination rate. Panels show the results for different values of the total selection st (increasing left to right) and ratio of migration probability to selection strength m/s (increasing top to bottom). Colours correspond to different values of the recombination rate asymmetry coefficient α: α = 1 (red), α =5 (green), α = 10 (cyan), α = 20 (purple). Error bars show 5th to 95th percentiles. Note that the highly stochastic nature in the results shown in the first column is caused by a very low number of realisations when a successful establishment occurred. Specifically, when the probability of establishment is close to 0 (see Fig.S9), there were only a few realisations from which this value could be calculated.*

### References

Venu, V., Harjunmaa, E., Dreau, A., Brady, S., Absher, D., Kingsley, D. M., & Jones, F. C. (2024). Fine-scale contemporary recombination variation and its fitness consequences in adaptively diverging stickleback fish. *Nature Ecology & Evolution*, *8*(7), 1337-1352. https://doi.org/10.1038/s41559-024-02434-4
